# Adipocyte-specific deletion of Dbc1 does not recapitulate healthy obesity phenotype but suggests regulation of inflammation signaling

**DOI:** 10.1101/2024.09.08.611743

**Authors:** Leonardo Santos, Rafael Sebastián Fort, Geraldine Schlapp, Karina Cal, Valentina Perez-Torrado, Maria Noel Meikle, Ana Paula Mulet, José R. Sotelo-Silveira, Jose M Verdes, Paola Contreras, Aldo J. Calliari, Martina Crispo, Jose L. Badano, Carlos Escande

## Abstract

The protein Deleted in Breast Cancer 1 (DBC1), a regulator of several transcription factors and epigenetic modulators, plays determinant roles in metabolism regulation, obesity and aging-related processes. Knockout mice for Dbc1, develop morbid obesity but are protected against liver steatosis, insulin resistance and atherosclerosis. We have proposed that this healthy obesity phenotype was mainly due to the expansion of adipose tissue, avoiding free-fatty acid spillover and metabolic damage in peripheral tissues. To gain more insight about the role of Dbc1 in adipose cells during obesity and its impact on metabolic dysregulation, we generated a conditional *Dbc1* KO mouse and backcrossed it with *CRE-AdipoQ* transgenic mice, aiming to abrogate Dbc1 expression in all mature adipocytes (cAT-Dbc1). cAT-Dbc1 mice showed deletion of Dbc1 specifically in mature adipocytes in different fat depots. We tested the effect of Dbc1 deletion in adipocytes on different aspects of metabolic regulation in male and female mice fed in normal chow and high-fat diets. We found that deletion of Dbc1 in mature adipocytes had no effect on weight gain, glucose tolerance and other markers of metabolic dysregulation, regardless sex. However, Dbc1 KO adipocytes displayed an mRNA expression profile consistent with increased inflammation during obesity. Our results suggest that the healthy phenotype displayed in the whole body Dbc1 KO obese mice is not due to the protein function in mature adipocytes and might involve other cell types present in adipose tissue. Instead, the specific deletion of Dbc1 in mature adipocytes highlights a novel role of Dbc1 in inflammation signaling during obesity.

## Introduction

Obesity and its related metabolic dysfunctions are a leading cause of morbimortality worldwide. While the irruption of GLP1 agonists is showing that treatment of obesity and allied comorbidities is no longer utopic, several aspects of the molecular signaling that determine the onset and progression of obesity-related metabolic dysfunction remain elusive. Accumulating evidence shows that the functional status of adipose tissue plays a determinant role on the onset and progression of systemic metabolic dysfunction during obesity. Adipose tissue inflammation, hypoxia and fatty-acid spillover are among the main determinants that affect metabolic dysfunction and determine the fate of several other organs, such as liver, pancreas and skeletal muscle (revised in [1]).

We and others have previously shown that the protein Deleted in Breast Cancer 1 (Dbc1) plays key roles in the onset and progression of metabolic dysfunction during obesity. We found that Dbc1 KO mice develop morbid obesity characterized by a healthy expansion of adipose tissue. Importantly, Dbc1 KO mice were protected against fat tissue inflammation and cellular senescence in adipose tissue [2], [3], fatty liver disease, and atherosclerosis [3].

Dbc1 was shown to bind and regulate several target proteins that are key players in metabolism regulation. Among them, there are transcription factors (p53, Foxp3, Rev-erbα), epigenetic modifiers, such as SIRT1, HDAC3 and SUV39H1, DNA repairing enzymes (BRCA1 and PARP1) and nuclear the receptors ER-α, ER-β. Interestingly, by binding and activating IKK-β at the cytoplasm, Dbc1 promotes NFkB-dependent transcriptional activation of various genes involved in inflammation and other immune responses[4], [5], [6], [7], [8], [9], [10].

Such promiscuity in its binding and regulation capacity, makes understanding the specific roles of Dbc1 a challenging issue. Indeed, the specific role of Dbc1 seems to depend on cell type and context. In an effort to understand the exact function of Dbc1 in regulating adipose tissue function during obesity, we engaged in generating a conditional, tissue-specific knockout model. We generated this mouse model by inserting LoxP sites the *Dbc1* gene using CRISPR/Cas9 technology. After obtaining the modified mouse, we backcrossed it with adipocyte-specific CRE transgenic mice (Tg(Adipoq-cre)1Evdr), in order to specifically delete Dbc1 in mature adipocytes. Adipocyte-specific Dbc1 KO males and females were characterized in normal chow and high-fat diets. We found that adipocyte-specific KO mice did not recapitulate our original findings in Dbc1 whole body KO mice. Both males and females gained similar weight and showed comparable results for all markers tested to control animals. Finally, we tested if deletion of Dbc1 in mature adipocytes had any functional consequence in gene expression profile during obesity. RNAseq data from adipocytes isolated from obese WT and conditional Dbc1 KO mice, showed a molecular fingerprint of inflammation in those animals where Dbc1 was deleted. The latter suggest that the healthy phenotype of the whole body Dbc1 KO mouse and assigned to a protection against adipose tissue inflammation, is independent of the role of Dbc1 in mature adipocytes. Alternatively, it may involve other cells present in the tissue. We conclude that in mature adipocytes, Dbc1 may still be playing a role in regulating inflammatory signaling during obesity, however the exact mechanism to achieve the so-called healthy phenotype seems to be different to that originally proposed.

## Material and methods

### Reagents and Antibodies

Unless otherwise specified, all reagents and chemicals were purchased from Sigma-Aldrich. Anti rabbit DBC1 antibody was purchased from Bethyl Laboratories (Cat. # A300-434A) and mouse monoclonal anti B-actin was from Sigma-Aldrich (Cat. # A2228).

### Animal handling and experiments

All mice used in this study were bred and maintained at the Institut Pasteur Montevideo Animal facility (UBAL). The experimental protocol was approved by the Institutional Animal Care and Use Committee of the Institut Pasteur Montevideo (CEUA; protocol numbers 70153-000839-17, 003- 19, and 006-19). All the studies described were performed according to the methods approved in the protocol and following all international guidelines and national legal regulations (Law 18.611). Mice received a standard chow or high-fat diet (42% fat and 0.25% cholesterol, AIN93G, LabDiet, USA) and water *ad libitum*.

### Generation of the Dbc1 conditional knock-out mice

All animal procedures to generate the mutant line were performed at the SPF animal facility of the Laboratory Animal Biotechnology Unit of Institut Pasteur de Montevideo. Experimental protocols were opportunely approved by the Institutional Animal Ethics Committee (protocol number 007-18), in accordance with national law 18.611 and international animal care guidelines (Guide for the Care and Use of Laboratory Animal) [17] regarding laboratory animal’s protocols. Mice were housed on individually ventilated cages (Tecniplast, Milan, Italy) containing chip bedding (Toplit 6, SAFE, Augy, France), in a controlled environment at 20 ± 1°C with a relative humidity of 40-60%, in a 14/10 h light-dark cycle. Autoclaved food (Labdiet 5K67, PMI Nutrition, IN, USA) and autoclaved filtered water were administered *ad libitum*.

Cytoplasmic microinjection was performed in B6D2F2 or C57BL/6J zygotes using a mix of 100 ng/µl Cas9 mRNA, 25 ng/µl of each sgRNA, and 50 ng/µl ssDNA oligo diluted in microinjection buffer. Viable embryos were cultured overnight in M16 medium microdrops under embryo tested mineral oil, containing ligase IV inhibitor SCR7, in 5% CO2 in air at 37 °C. The next day, 2-cell embryos were transferred into the oviduct of B6D2F1 0.5 days poscoitum (dpc) pseudopregnant females (20 embryos/female in average), following surgical procedures established in the animal facility [18]. For surgery, recipient females were anesthetized with a mixture of ketamine (100 mg/kg, Pharmaservice, Ripoll Vet, Montevideo, Uruguay) and xylazine (10 mg/kg, Seton 2%; Calier, Montevideo, Uruguay). Tolfenamic acid was administered subcutaneously (1 mg/kg, Tolfedine, Vetoquinol, Madrid, Spain) in order to provide analgesia and anti-inflammatory effects [19]. Pregnancy diagnosis was determined by visual inspection by an experienced animal caretaker two weeks after embryo transfer, and litter size was recorded on day 7 after birth.

Pups were tail-biopsied and genotyped 21 days after birth, and mutant animals were maintained as founders. First round of microinjections resulted in the insertion of the 3’ LoxP site only. *Dbc1* 3’ LoxP F1 animals were used to produce embryos by *in vitro* fertilization using CARD (Center for Animal Resources and Development) protocol [20]. Three hours after fertilization, zygotes were microinjected into the cytoplasm with CRISPR reagents in the same aforementioned concentrations, in order to insert the lacking 5’ LoxP site. Two-cell viable embryos were transferred to pseudopregnant females as detailed above. Founders with 3’-5’ LoxP sites were used to generate the final mutant line.

The mice used in this study were backcrossed with C57JB6 mice to homogenize the genetic background. They were then bred to homozygosity for the floxed Dbc1 allele. To generate the adipocyte-specific DBC1 knockout mice (AT Dbc1 KO), the homozygous floxed DBC1 mice were crossed with B6.FVB-Tg(Adipoq-cre)1Evdr/J mice obtained from Jackson Laboratories.

The experimental colony was established by crossing Adipoq-cre Dbc1^LoxP/LoxP^ hemizygous mice with Dbc1^LoxP/LoxP^ mice. In the experiments, groups with a sample size (N) of ten or more mice per genotype were used.

### Blood glucose measurements

Mice were kept in fasting for 16 h before fasting glucose measurement and glucose tolerance tests (GTT). For GTT, mice were injected (IP) with 1.5 g/kg body weight of glucose solution. Plasma glucose concentrations were measured from blood obtained from the tail using a hand- held glucometer (Accu-Chek, Roche).

### Hepatic and renal function

To assess liver and kidney function, whole blood from cardiac puncture was analyzed using the Pointcare V2 Chemistry Analyzer (Tianjin MNCHIP Technologies Co., China). The parameters determined were total proteins (TP), albumin (ALB) level, globulin (GLO) level, ALB/GLO ratio, total bilirubin (TBIL) level, alanine aminotransferase (ALT) level, aspartate aminotransferase (AST) level, gamma glutamyl transpeptidase (GGT) level, blood urea nitrogen (BUN) level, creatinine (CRE) level, BUN/CRE ratio and glucose (GLU) level.

### Isolation of WAT adipocytes

Fat depots were dissected and weighed. Collagenase digestion solution (collagenase type II, GIBCO 1 mg/ml in Hank’s Balanced Salt Solution containing 2% BSA) was added in 3:1 collagenase fat tissue ratio, and tissue was fined minced and incubated for 1 hour at 37°C in a shaking incubator. Then adipocyte suspension was centrifuged at 100g, 10 minutes at RT. Subsequently, floating mature adipocytes were washed twice with cold PBS.

### Western blotting

Adipocytes were lysed using radioimmunoprecipitation assay buffer (in a volume ratio of 1:10) supplemented with 5 mM NaF, 5 mM nicotinamide, 50 mM β-glycerophosphate, 1 μM trichostatin A (catalog no.: 647925; Sigma), a protease inhibitor cocktail, and then sonicated by means of five cycles of 10 s each followed by intervals of 5 s. Homogenates were incubated during 20 to 30 min at 4 °C under constant agitation and then centrifuged at 10,000*g* during 10 min. Protein concentrations in the supernatants were determined using the Bradford protein assay reagent. Samples were resuspended in Laemmli 5×, separated in SDS-PAGE gels, and transferred to polyvinylidene fluoride membranes. After blocking (with Tris-buffered saline containing 0.2% Tween-20 and 5% nonfat milk), the membranes were incubated overnight with the appropriate antibodies. Secondary antibodies were incubated 1 h and detected using SuperSignal West Pico Chemiluminescent Kit (catalog no.: 34080; Pierce).

### Histology

Liver sections were obtained from paraffin-embedded tissue and stained with H&E following standard procedures. For each mouse, at least 3 sections were analyzed.

### Plasma NEFA quantification

Blood was drawn from mice (by cardiac puncture using deep anesthesia). Non-esterified fatty acids were measured by using NEFA-HR Assay (NEFA,, Waco) according to the manufacturer’s instructions.

### Lipids accumulation in liver measurements

Liver Triglycerides was measured according to[11]. Samples were homogenized in the lysis buffer as described above. Lipid content was measured by using the TG Color GPO/PAP AA (Wiener Lab, USA) according to the manufacturer’s instructions.

### Adipocyte Lysis and RNA Extraction

Total RNA from adipocytes was extracted following lysis with Trizol reagent (Thermo Fisher Scientific) according to the manufacturer’s instructions. Briefly, adipocytes were harvested and resuspended in 1 mL of Trizol reagent. The cells were lysed by pipetting up and down several times and then incubated at room temperature for 5 minutes to ensure complete dissociation of nucleoprotein complexes. Following lysis, 0.2 mL of chloroform was added per 1 mL of Trizol reagent. The mixture was vigorously shaken for 15 seconds and then incubated at room temperature for 2-3 minutes. The samples were centrifuged at 12,000 x g for 15 minutes at 4°C. The aqueous phase was then processed using the Direct-zol RNA Miniprep Kit (Zymo Research) according to the manufacturer’s protocol.

### RNA quality control, library and sequencing

An Infinite 200 PRO (Tecan) spectrophotometer was used to evaluate the concentration and the 260/280 ratios of RNA obtained from mature adipocytes.The RNA quality was evaluated using the RNA integrity number (RIN), determined with a Bioanalyzer RNA 6000 Nano chip (Agilent, Cat# 5067-1511). All samples demonstrated high RNA integrity, with an average RIN of 9.0 were used for further studies. RNA-seq was performed on samples from individual animals (n ≥ 3) of each experimental group. All libraries were prepared and sequenced by Genewiz (https://www.genewiz.com/) using the TruSeq mRNA Sample Prep Kit v2 protocol (poly(A)+ selection) and Illumina HiSeq 2x150 bp sequencing. The raw FASTQ data sets supporting the results of this article are available at the Sequence Read Archive repository (https://www.ncbi.nlm.nih.gov/sra) (BioProject: PRJNA1157834)

### RNA-seq analysis

Raw reads (Fastq files) obtained from Genewiz (https://www.genewiz.com/) were used as the input for miARma-Seq pipeline, a comprehensive tool for miRNA and mRNA analysis [21], following the user manual (v 1.7.2). Low quality reads and adapter sequences were removed with Cutadapt software [22] allowing a minimum read Phred quality of 20. Filtered high quality reads (Ave. 54,995,205 ± SD 6,123,659 reads, Supplementary Table 1) were aligned to Mus musculus reference genome (GRCm38/mm10 indexed from http://bowtie-bio.sourceforge.net/bowtie2/index.shtml) using Hisat2 [23] with default parameters (Overall genome mapping %: Ave. 94.9 ± SD 0.3, Supplementary Table 1). Gene counts were assessed with featureCounts [24]using default parameters (Supplementary Table 1) For quantification we use the annotation coordinates of ensembl Mus_musculus.GRCm38.96 GTF file. Normalization and Differential Expression Genes (DEGs) analysis were conducted using SARTools pipeline [25] in RStudio, selecting edgeR algorithm [26], Trimmed Mean of the M-values (TMM) normalization and CPM (Counts Per Million) ≥ 1 as cut-off (Supplementary Table 1 and Supplementary Figure 4). A significance threshold of log2 fold change > |0.58| and p-value <0.05 was applied for DEGs, instead of adjusted p-value, in order to have a sufficient number of genes to perform over- representation analysis.

### Gene Set Enrichment Analysis

Gene Set Enrichment Analysis (GSEA) was performed using gseGO (R package clusterProfiler v4.8.2) [14] to explore the Gene Ontology (GO) terms (Biological Process) using the complete list of genes (ranked based on its fold change of expression), extending beyond the analysis of DEGs. The terms with significance threshold of adjusted p-value <0.05 were selected.

### Statistics

Values are presented as mean ± SD of three to five experiments, unless otherwise indicated. The significance of differences between means was assessed by ANOVA or a two-tailed Student *t* test, as indicated. R software (version 3.6) was used for plots of RNA-seq data using libraries: enrichplot, xlsx, ggplot2, heatmap.2 (clustering distance measured by Euclidean and Ward clustering algorithms). A *P* value of <0.05 was considered significant.

### Data availability

The datasets supporting the conclusions of this article are available at SRA repository (BioProject: PRJNA1157834) and within the article’s supplementary information.

## Results

### Generation of Dbc1^LoxP/LoxP^ conditional mice by CRISPR/Cas9 and adipocyte-specific deletion

The mouse *Dbc1* gene, located on chromosome 14, comprises 21 exons forming an open reading frame that extends from exon 2 to exon 21. It is predicted to have only one full-length transcript encoding a protein of 922 amino acids. To create a conditional knockout (cKO) murine line, we targeted exon 3 to be floxed. We designed two guide RNAs (gRNAs) to target intron 2 and 3 using CRISPR/Cas9 and assessed their effectiveness by transiently transfecting a murine NIH3T3 cell line that stably expresses Cas9, followed by heteroduplex analysis (Fig. S1).

Additionally, we generated two single-stranded DNA (ssDNA) donors for knock-in loxP sites. This sequence comprises part of intron 2 and 3, LoxP sites, and two restriction enzyme sites, which allow for the identification of the insertion. In the first step, we achieved the insertion of the 3’ loxP site. A second round of genetic manipulation was necessary to insert the 5’ loxP site.

We crossed a male mouse carrying the selected mutation with C57BL/6J females to establish the colony and performed five rounds of backcrossing before beginning the characterization of the line. This process should be sufficient to segregate any off-target or unwanted modifications of the genome produced during genetic manipulation. From that point onward, we maintained the line by crossing homozygous mutant animals. To achieve the knock-out in mature adipocytes, we crossed the floxed *Dbc1* animals with the *AdipoQ-CRE* mouse line. (Fig 1A).

**Figure 1.**
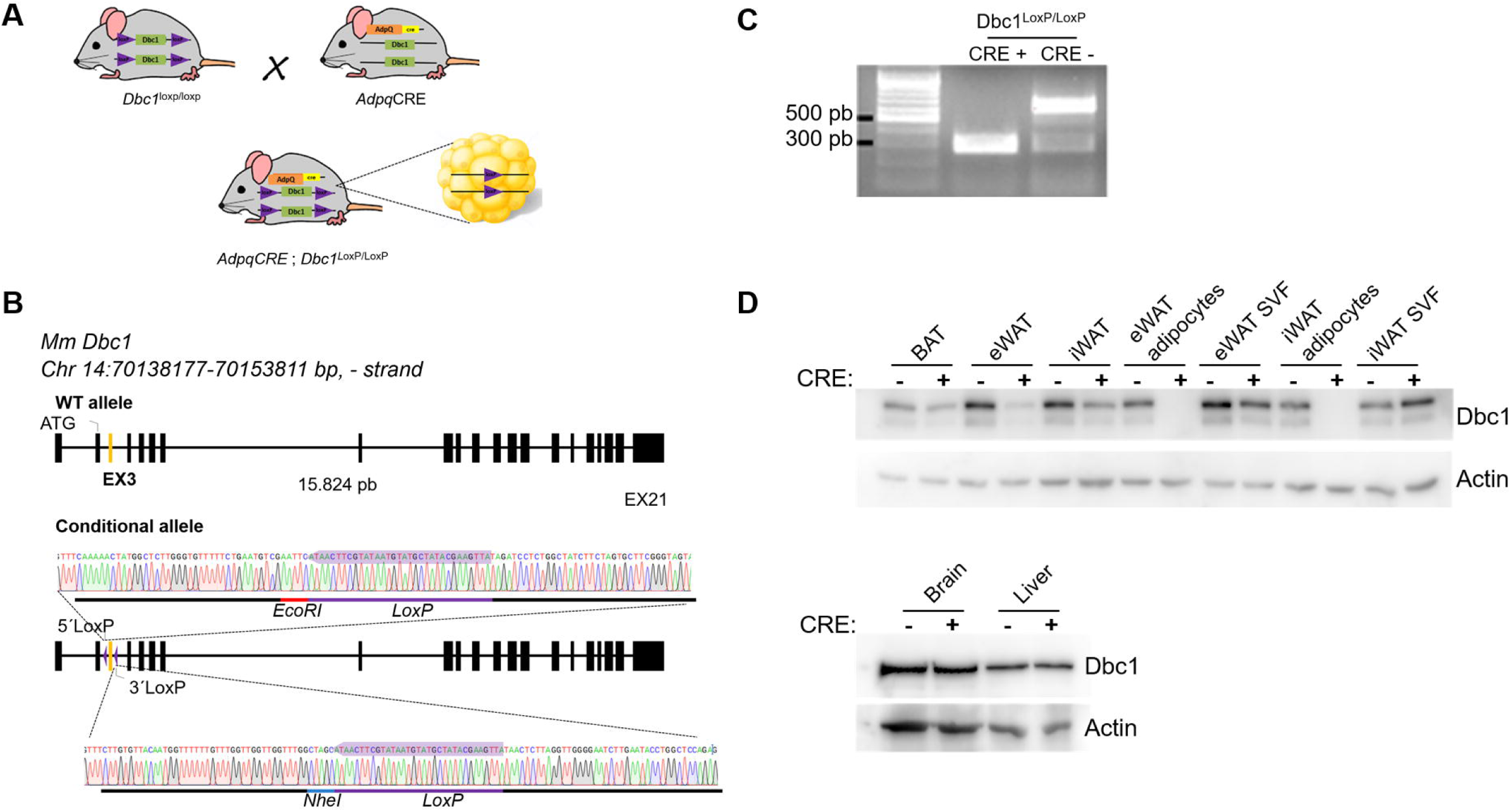
Generation of Adipocyte-Specific Dbc1 Knockout Mouse Model. **(A)** Breeding Strategy for Adipocyte-Specific Dbc1 Knockout. The use of *AdpqCRE* transgene in combination with *Dbc1* floxed allele resulted in the adipocyte-specific Dbc1 gene edition (**B)** Schematic Representation of LoxP Sites Inserted Flanking Exon 3 of *Dbc1* Gene.Illustration depicting the precise location of LoxP sites inserted around exon 3 of the Dbc1 gene. **(C)** PCR Detection of Adipocyte-Specific Dbc1 Gene Edition. PCR analysis of the offspring carrying both the Dbc1 floxed allele and *AdpqCRE* transgene, demonstrating the specific Dbc1 gene edition in adipocytes. **(D)** Western Blot Analysis of Dbc1 Protein Expression. Images of Western blot analysis showcasing the expression of Dbc1 protein in inguinal adipose tissue (iWAT), epididymal adipose tissue (eWAT), isolated adipocytes, stromal vascular fraction, brain, and liver.

As a first step, we sequenced the region where we made the genetic manipulation and confirmed that the sequence was as expected. Using primers located at the nearest 5’ and 3’ ends of Dbc1 exon 3, we sequenced the PCR product. We detected the expected mutations and found no evidence of unwanted mutations (Fig. 1B). To verify the construct, we performed PCR on genomic DNA from cKO MEF cells transiently expressing Cre recombinase. A 300 bp PCR product indicated successful DNA recombination (Fig 1C).

Next, we evaluated Dbc1 at the protein level by western blot. The ablation of exon 3 (300 pb) by Cre recombinase was expected to cause a shift in the open reading frame of the transcript and the introduction of a premature stop codon. The mutant allele could therefore produce a 35-amino acid peptide. In agreement with that prediction, the ∼100 kDa Dbc1 band was absent in adipocytes from different fat tissue depots. Also, in whole tissue extracts from white adipose tissue (WAT) and brown adipose tissue (BAT), Dbc1 protein levels decreased. In contrast, there was no difference in protein expression in the liver and brain (Fig. 1D).

### Metabolic characterization of cAT-Dbc1 KO male and female mice on normal diet

In previous studies, we and others showed that whole-body Dbc1 knockout mice do not develop obesity when fed a regular chow diet [11], [12]. However, they present abnormal glucose management. We showed that this effect was mainly due to altered liver gluconeogenesis [12]. This effect was a consequence of dysregulation of hepatic rev-erb alpha activity and increased PEPCK expression [11], [12]. Consistent with a specific deletion only in mature adipocytes, we found that cAT-Dbc1 KO mice had no affectation of body weight or glucose management, measured by GTT in normal chow conditions, both in male and female mice (Figure 2). Free-fatty acid levels were also comparable among genotypes both in males and females (Figure 2). In conclusion, the absence of Dbc1 specifically in mature adipocytes failed to influence body weight, glucose management, or free-fatty acid levels in mice under normal chow conditions.

**Figure 2.**
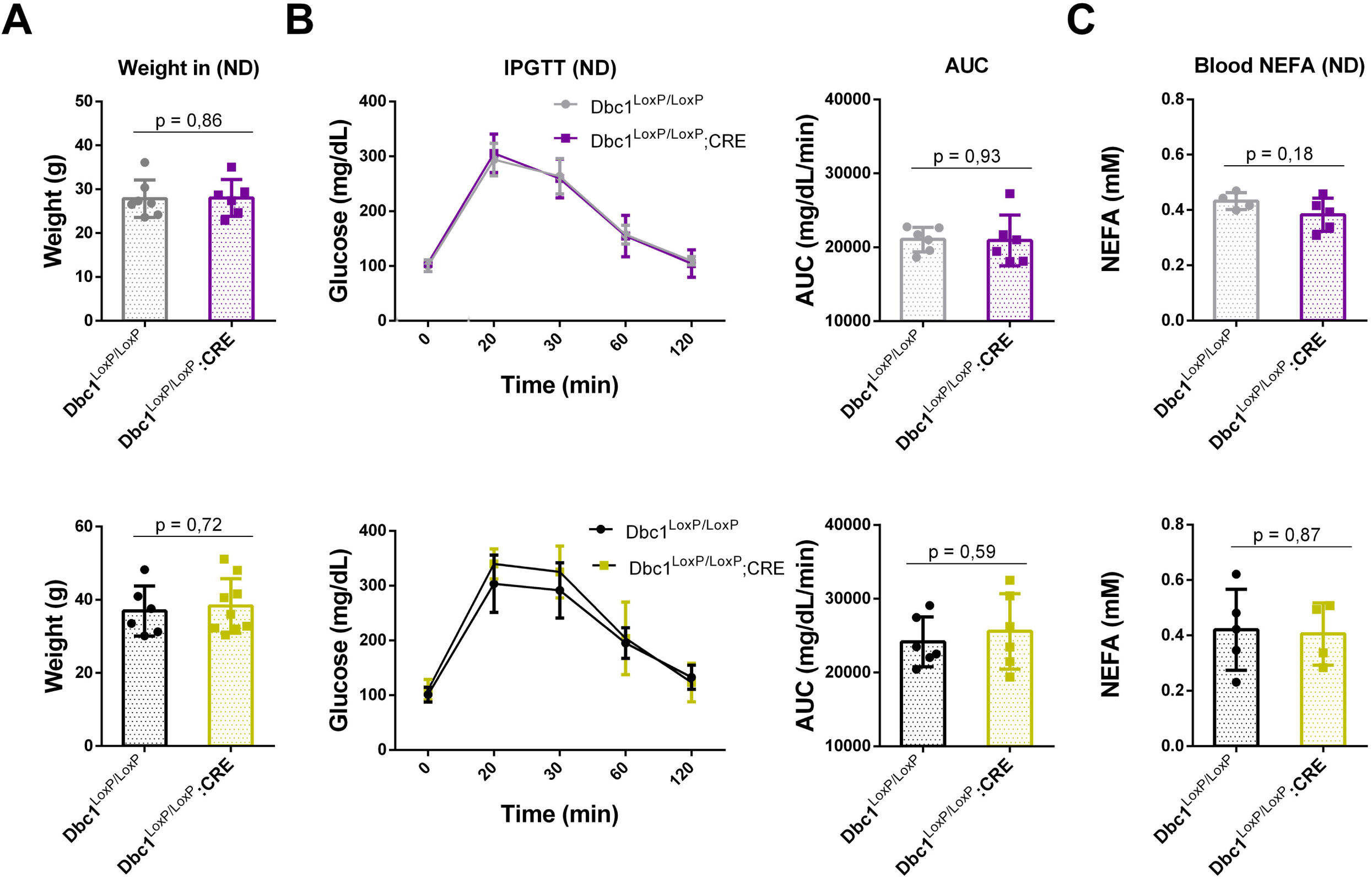
Similar Weight, Glucose, and Blood Free Fatty Acid Levels Across Different Genotypes under Normal Diet (ND). **(A)** Mean body weight of female (gray and purple) and male mice (black and yellow) from different genotypes **(B)** Glucose Tolerance Test (GTT). Glucose response evaluation in male and female mice from different genotypes **(C)** Blood Free Fatty Acid Levels of male and female mice from different genotypes. Data represent the mean ± standard deviation (SD) for each group. Statistical analysis was performed using the t-test, and no statistically significant differences were observed.

### Metabolic characterization of cAT-Dbc1 KO male and female mice on high-fat diet

Next, we evaluated the effect of diet-induced obesity on the metabolic phenotype of cAT-Dbc1 KO mice. Our previous work, in mice with mixed genetic background, showed that whole-body Dbc1 KO mice develop morbid obesity with prevention of fatty-acid spillover and metabolic damage [11]. This phenotype was recapitulated in mice with pure C57BL/6J genetic background (Supplementary Figure 2). However, when we fed cAT-Dbc1 with a high-fat diet (HFD), we found that both males and females gained similar weight than their control littermates (Figure 3A). We followed weight gain for up to 20 weeks of a high-fat diet with no difference among genotypes.

**Figure 3.**
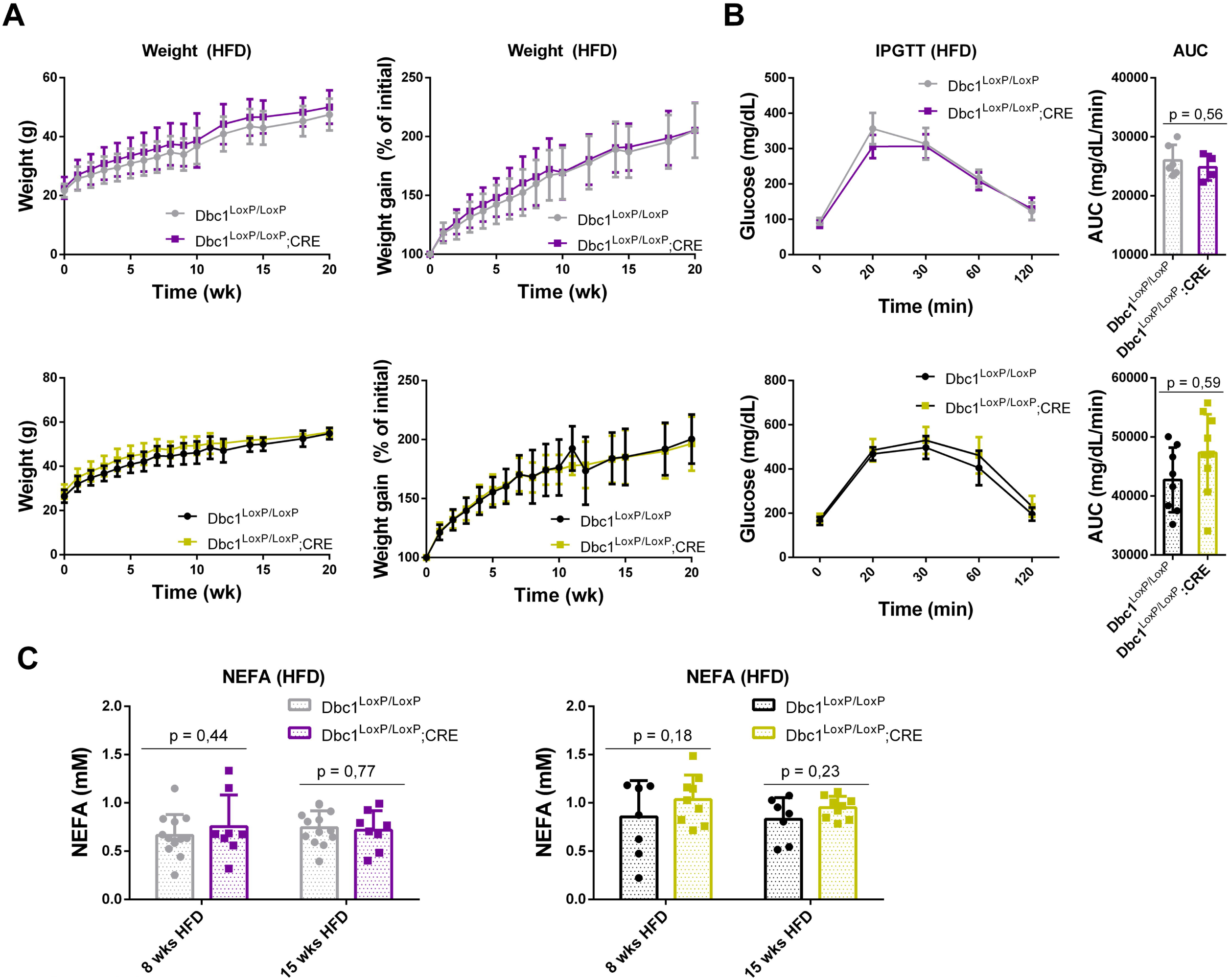
Development of the Diet Induced Obesity (DIO) in cAT-Dbc1 mouse model (A) Comparable weight gain was observed across both sexes and genotypes during HFD administration, with no significant differences detected. (B) Glucose tolerance test was assessed after 8 weeks on the HFD regimen. No significant differences were found in glucose tolerance between the experimental groups. (C) Plasma Free Fatty Acid Levels of serum samples collected after 8 and 15 weeks of HFD administration were analyzed for free fatty acid

Conversely, we did not observe any differences in glucose management (Figure 3B), or in levels of free-fatty acids in plasma (Figure 3C), the hallmark of the healthy obesity phenotype that we described before [3].

We measured several markers of liver and renal function after 15 weeks of a high-fat diet, both in males and females. Despite a clear effect of diet upon the levels of several markers, we found no differences among genotypes (Figure 4). Kidney function markers such as the A/G ratio and BUN/CRE indicate that renal function was normal compared to levels in chow-fed mice. Hepatic enzymes alanine aminotransferase (ALT) and aspartate aminotransferase (AST) show a significant increase in blood levels compared to lean mice, indicating an effect of the high-fat diet on hepatic physiology. However, bilirubin levels and gamma glutamyl transferase (GGT) activity did not change relative to controls, suggesting that liver damage was moderate. Under HFD, both genotypes accumulate triglycerides in the liver without significant differences.

**Figure 4.**
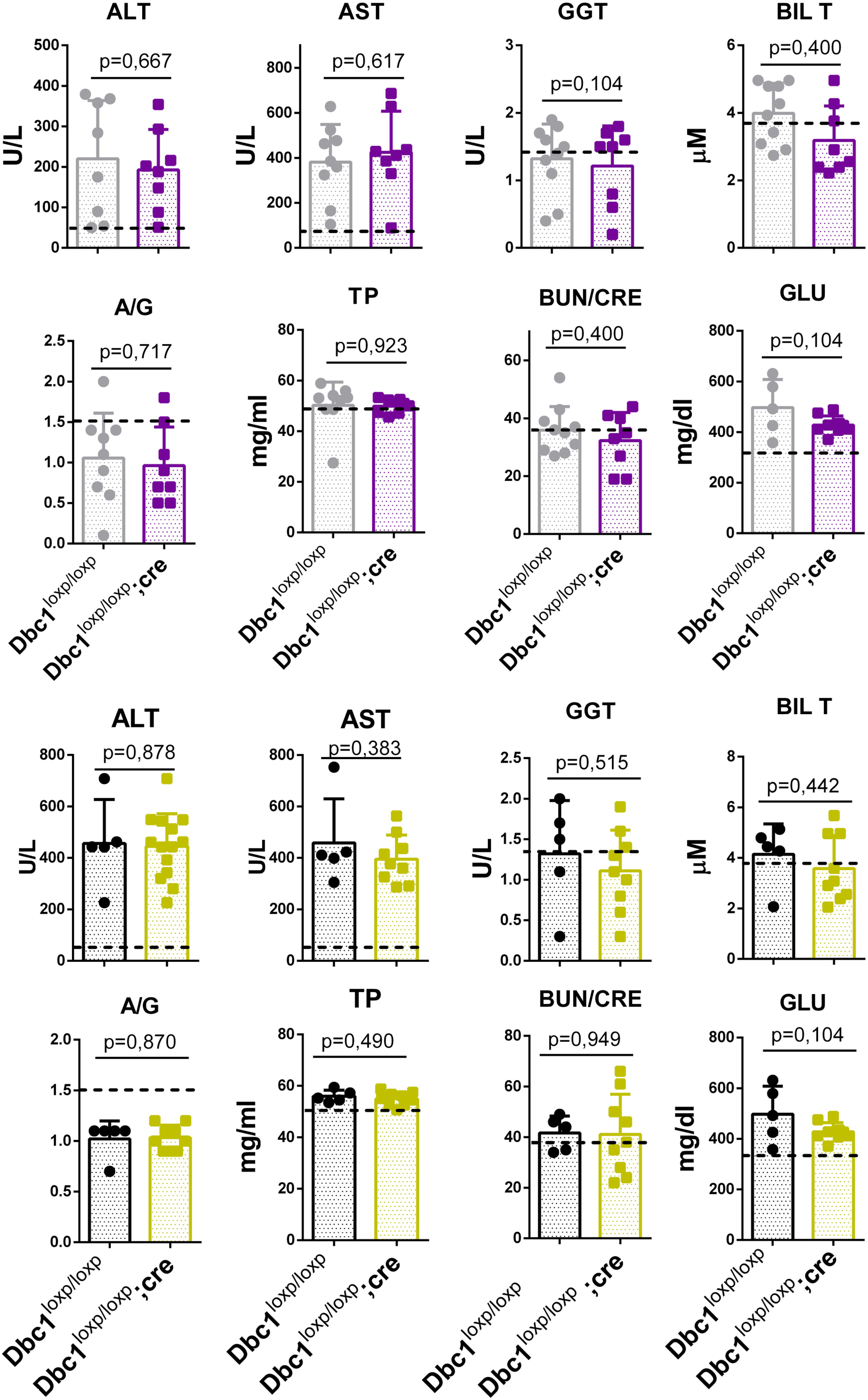
Renal and Hepatic Function in Male and Female Mice of Both Genotypes. Graphs illustrating liver and kidney function data obtained from blood analysis of female (gray and purple) and male mice (black and yellow) following 15 weeks of high-fat diet (HFD) administration. The data represent the evaluation of renal and hepatic markers, including total bilirubin (TBIL), blood urea nitrogen to creatinine ratio (BUN/CRE), total protein (TP), albumin and globulin levels (ALB and GLOB), aspartate aminotransferase (AST), gamma-glutamyl transferase (GGT), alanine transaminase (ALT), albumin-to-globulin ratio (A/G), and non-fasting glucose (GLU). The dashed line shows biomarker levels from normal chow-fed mice. Results are shown for both genotypes. Data represent the mean ± standard deviation (SD) for each group. Statistical analysis was performed using the t-test.

Histological analysis of liver samples also showed steatosis, as is expected in the diet induced obesity (DIO) condition (Figure 5). Overall, cAT-Dbc1 KO mice show no significant differences in weight gain, glucose management, or liver and kidney function compared to controls under a high-fat diet treatment, indicating that adipocyte-specific Dbc1 deletion does not replicate the protective effects observed in whole-body Dbc1 knockout models. levels, with no significant differences detected between genotypes. Data represent mean ± standard deviation (SD).

**Figure 5.**
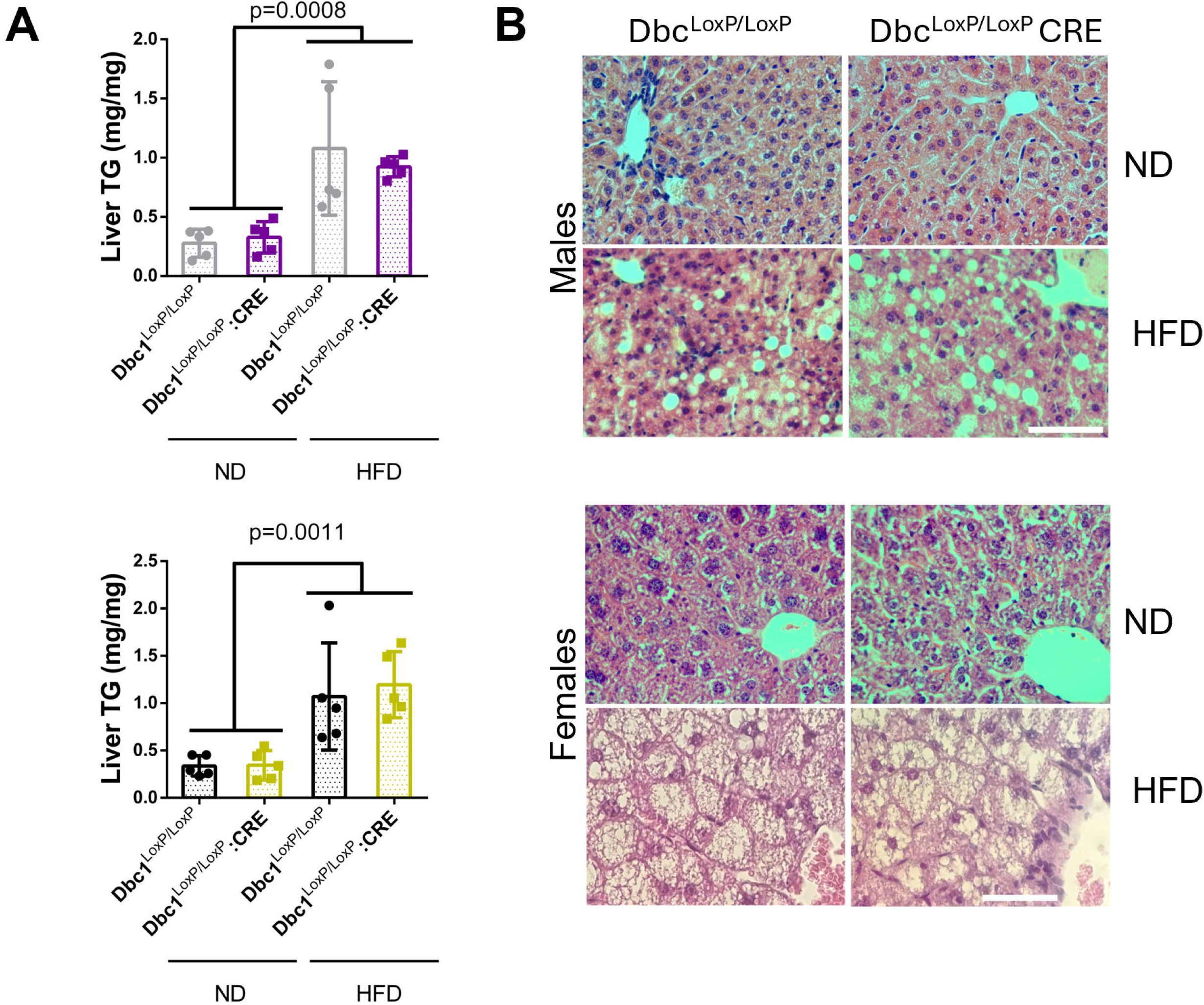
Hepatic Responses to Diet induced Obesity. **(A)** Liver Triglyceride Content in Both sex and Genotypes under Normal Diet and High-Fat Diet (HFD). Data represent mean ± standard deviation (SD). Statistical analysis was performed using two-way ANOVA (**B)** Hematoxylin and Eosin Histology of Male Livers under Normal Diet and High-Fat Diet. Scale Bar, 50 μm.

### RNA-seq analysis from isolated white adipocytes in WT and cAT-Dbc1 KO obese mice

Finally, in an effort to determine the role Dbc1 is playing in mature adipocytes during obesity, we performed RNA-seq analysis of isolated adipocytes from cAT-Dbc1 and their littermates control after 20 weeks of high-fat diet. Whole transcriptome sequencing was conducted on five samples of adipocytes from cAT-Dbc1 knockout mice and three control mice. We performed transcriptome analyses to identify differentially expressed genes (DEGs) and functional pathways using bioinformatics methods. RNA-Seq revealed 60 DEGs with log2FC |0.58| and p-value < 0.05 between control and Dbc1 KO adipocytes (Figure 6A). Among these, 42 genes were upregulated in Dbc1 KO adipocytes, while 18 genes were downregulated in control adipocytes with a 1.5-fold change (Figure 6B). Overall, the data indicate that Dbc1 moderately influences the transcriptional landscape of adipocytes during diet-induced obesity, potentially affecting their function. To explore the functions of the identified DEGs list, we analyzed which pathways and cellular processes were enriched for each comparison using Enrichr (https://maayanlab.cloud/Enrichr/ and the database Reactome 2022) [13]. We found significant pathways enriched only for the list of upregulated DEGs, the top three pathways are related to immune system response: Immune System, Cytokine Signalling in Immune System, and Signalling of Interleukins (Figure 6C). To surpass the constraints of over-representation analysis based on a limited number of DEGs, we decided to use Gene Set Enrichment Analysis (GSEA) involving the entire set of genes in the transcriptome. This is a well-known strategy that permits exploring the complete list of genes ranked based on the FC of expression between conditions [14]. We conduct this analysis for Gene Ontology terms using the ClusterProfiler algorithm. This approach identified significantly enriched Gene Ontology terms (Biological Process). According to the GO dataset, Dbc1 knockout (KO) adipocytes showed global upregulation in several inflammatory response pathways (Figure 6D). Particularly noticeable are the positive enriched terms related with cytokines production and NFkB transcriptional activity (Figure 6E). Globally, this analysis underscores the involvement of DBC1 in regulating inflammatory responses in adipocytes.

**Figure 6:**
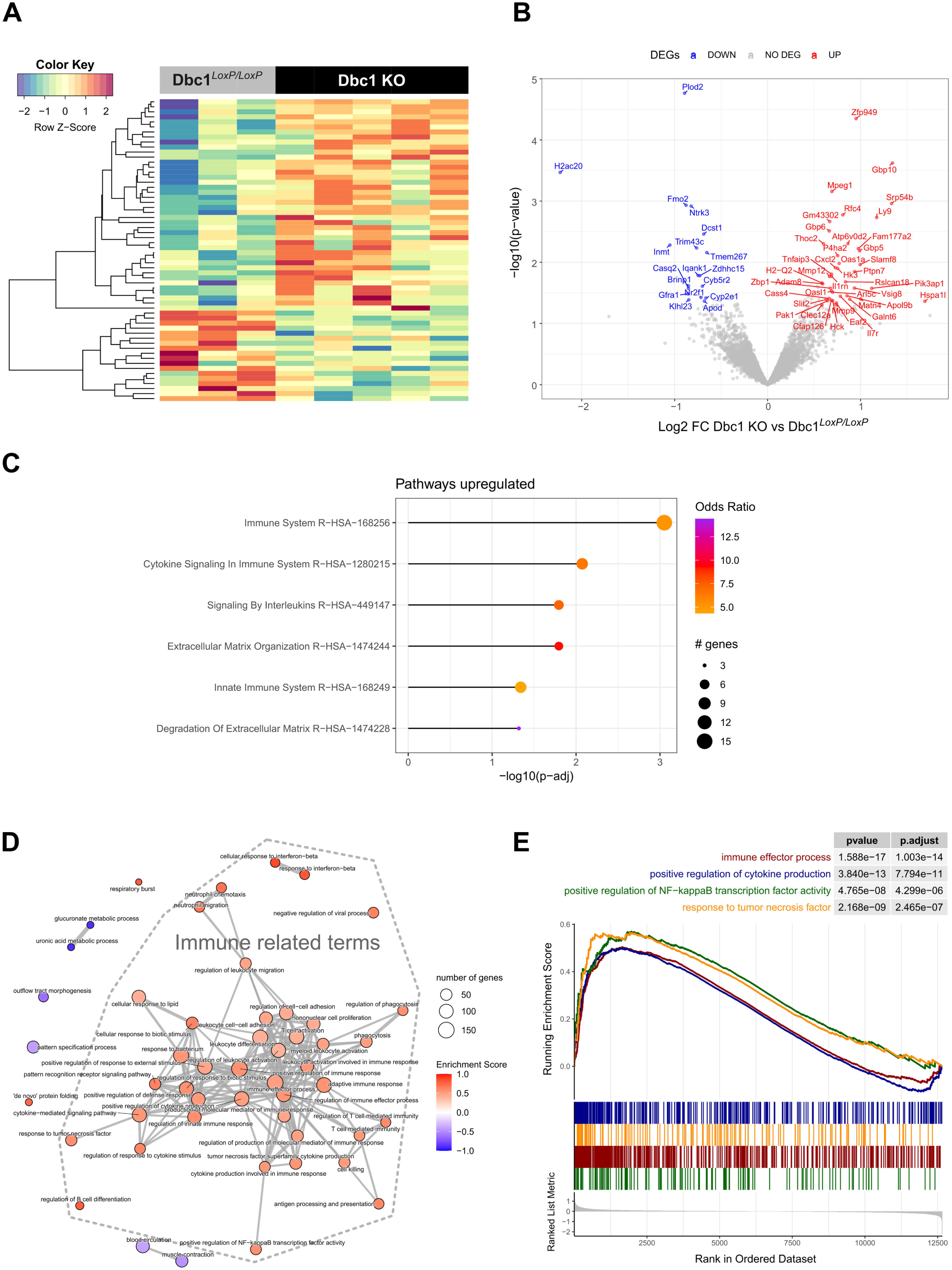
Transcriptomic data of mature adipocytes Dbc1 KO vs WT. **(A)** Heatmap of differentially expressed protein coding genes (DEGs) with log2FC |0.58| and p-value < 0.05. **(B)** Volcano plot of DEGs in red and blue are shown upregulated and downregulated DEGs, respectively. **(C)** Reactome enriched pathways are shown for upregulated DEGs. **(D)** Emapplot of the top 50 significantly enriched Gene Ontology terms (Biological Process) (increased in red or decreased blue in the Dbc1 KO compared to the Dbc1^Lxp/Loxp^. **(E)** Gene Set Enrichment Analysis (GSEA) of Gene Ontology terms selected from **(C)** The x-axis represents the rank for all genes based on log2 (Fold Change); the y-axis (upper panel) shown the Enrichment Score value, the y-axis (bottom panel) represents the value log2FC of the ranking metric.

## Discussion

As a regulator of several transcription factors and epigenetic modulators, Dbc1 has emerged as a multifaceted determinant in metabolism regulation. Interestingly, Dbc1 knockout mice develop morbid obesity but are protected against liver steatosis, insulin resistance and atherosclerosis. It has been proposed that this expansion in the adipose tissue avoids free-fatty acid spillover and metabolic damage in peripheral tissues, thus accounting for the observed healthy phenotype. In order to further understand the roles of Dbc1 in adipose tissue and its associated phenotype, we generated a conditional knockout mouse model aiming to specifically delete Dbc1 in mature adipocytes.

The generation of Dbc1 conditional KO mice was successful, confirmed by the correct insertion of LoxP sites and the absence of unwanted mutations. Adipocyte-specific deletion of Dbc1 was validated through PCR and western blot, showing the ablation of Dbc1 protein in adipocytes from cAT-Dbc1 KO animals. Western blot analysis of whole adipose tissue samples revealed that, in addition to adipocytes, the stromal vascular fraction also expresses high levels of the protein. This precise genetic manipulation enabled us to isolate the role of Dbc1 specifically in adipocytes, independent of its functions in other tissues.

Our metabolic characterization of cAT-Dbc1 KO mice under normal diet conditions showed no significant differences in body weight, glucose tolerance, or free-fatty acid levels compared to control mice. Under high-fat diet conditions, both male and female cAT-Dbc1 KO mice gained weight similarly to their control littermates, with no significant differences in glucose management, free-fatty acid levels, or markers of liver and renal function. These observations contrast with previous studies on whole-body Dbc1 knockout mice, which exhibited altered glucose management due to liver-specific mechanisms.

The failure to recapitulate the whole-body Dbc1 knockout phenotype in cAT-Dbc1 KO mice, indicates that the Dbc1 of adipocytes is not responsible for the overall metabolic health or susceptibility to diet-induced obesity. Instead, Dbc1 expressed in adipocytes might play a more subtle or context-dependent role, a possibility that requires further investigation under specific stress or metabolic conditions. As a whole, these findings put forward that the metabolic protection observed in whole-body Dbc1 knockout models may arise from non-adipocyte cells located into or even outside the adipose tissue. Moreover, the previous observed healthy obesity phenotype could be a result of complex interactions between different organs rather than being specific to a particular tissue or cell type. Further investigation is needed to clarify these results.

Our RNA-Seq analysis provided new insights into the molecular functions of Dbc1 in adipocytes. We identified several inflammatory pathways that are overrepresented in Dbc1 KO adipocytes. Notably, there is evidence of an inflammatory repressor role for Dbc1 in lymphocytes [15]. In fact, Dbc1 selectively suppresses alternative NF-κB pathways, which is in line with our GSA analisis (Fig supplementary 3). These findings are paradoxical, and intriguing given the diminished inflammation observed in the adipose tissue of whole-body Dbc1 KO mice [3]. Interestingly, it has been reported that adipocyte inflammatory pathways, such as TNF signaling, are important for proper adipogenesis [16]. Therefore, it could be hypothesized that a reduced ability to sense and respond to proinflammatory stimuli at the level of the adipocyte leads to a decrease in the capacity for healthy adipose tissue expansion and remodeling Thus, the elevated immune response in Dbc1 KO adipocytes observed in this study could be biologically relevant for the healthy expansion of adipose tissue seen in the whole-body KO model.

The upregulation of inflammatory pathways observed in Dbc1 knockout adipocytes points to a complex role for Dbc1 in maintaining adipocyte homeostasis and makes it difficult to give a simplistic explanation that reconciles with the diminished adipose tissue inflammation whole-body Dbc1 KO mice observed before [3]. Clearly, other cellular components of adipose tissue cooperate in maintaining adipose tissue homeostasis. The mechanistic links between Dbc1 and these inflammatory processes warrant further exploration, particularly to understand whether Dbc1 directly interacts with key signaling molecules involved in these pathways.

Our study challenges the belief that adipocyte-expressed Dbc1 protects against liver steatosis, insulin resistance, and atherosclerosis, instead revealing a nuanced role of Dbc1 in regulating gene expression and inflammatory responses in adipocytes, particularly under diet-induced obesity. Despite the lack of significant metabolic alterations following adipocyte-specific Dbc1 deletion, the observed transcriptional changes suggest that Dbc1 may be crucial for adipocyte function during obesity.

## Acknowledgements

CE was supported by grants by ANII (FCE_1_2014_1_104002), CSIC, PEDECIBA and FOCEM (COF 03/11). AC was supported by the CSIC-DT program, from the Udelar. LS was supported by grant FVF_2023_504. This work was supported by FOCEM - Fondo para la Convergencia Estructural del Mercosur (COF 03/11).

## Supporting Information

**Figure S1.**
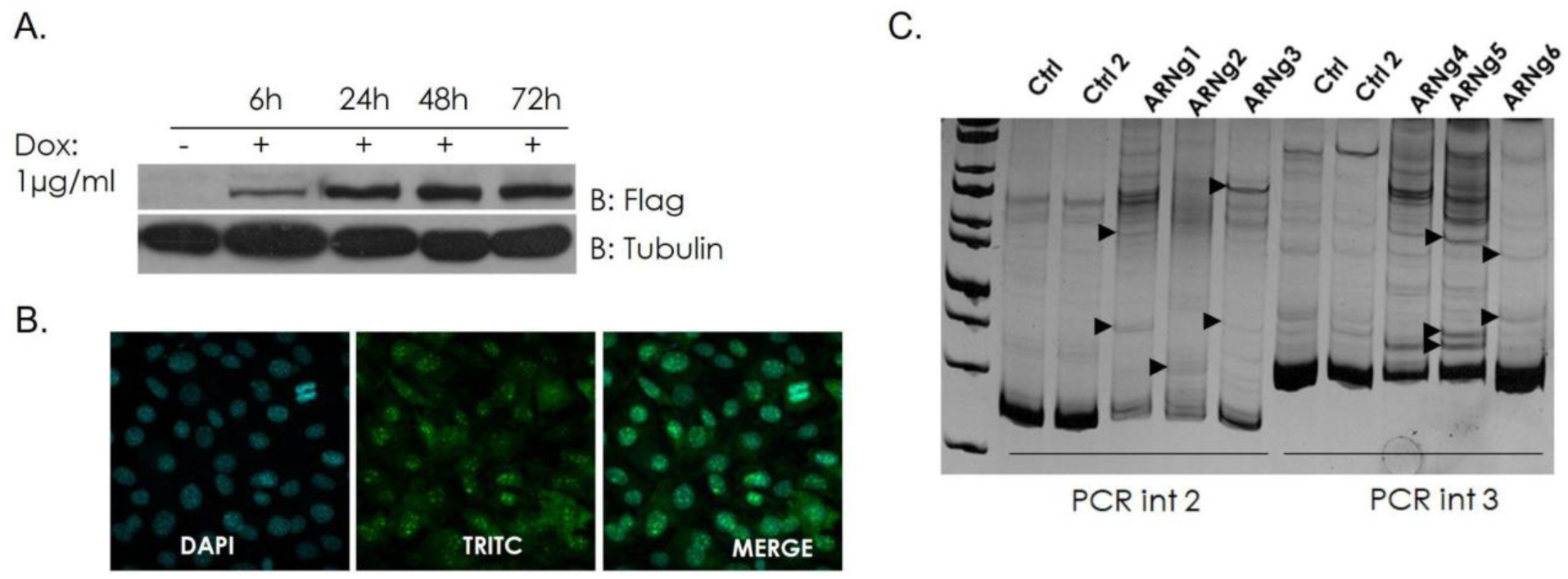
Testing of gRNA Efficiency in Inducible 3T3-Cas9 Cells. **(A)** Western blot analysis of total cell extracts at various time points following incubation with doxycycline. We generated a 3T3 L1 cell line where Cas9 expression is controlled by doxycycline. **(B)** Immunofluorescence images of cells after 48 hours of incubation with doxycycline. Nuclei are stained with DAPI, and Flag-Cas9 is detected by immunofluorescence following doxycycline induction. **(C)** Acrylamide gel electrophoresis stained with EtBr showing PCR products using genomic DNA from 3T3-Cas9 cells after transfection with various gRNAs and treatment with doxycycline. Black arrows indicate the formation of heteroduplexes, which are absent in the controls of uninduced and untransfected cells, as well as in induced but untransfected cells (Ctrl and Ctrl2, respectively). A noticeable decrease in amplicon yield suggests significant modifications following Cas9 cleavage and DNA repair processes.

**Figure S2.**
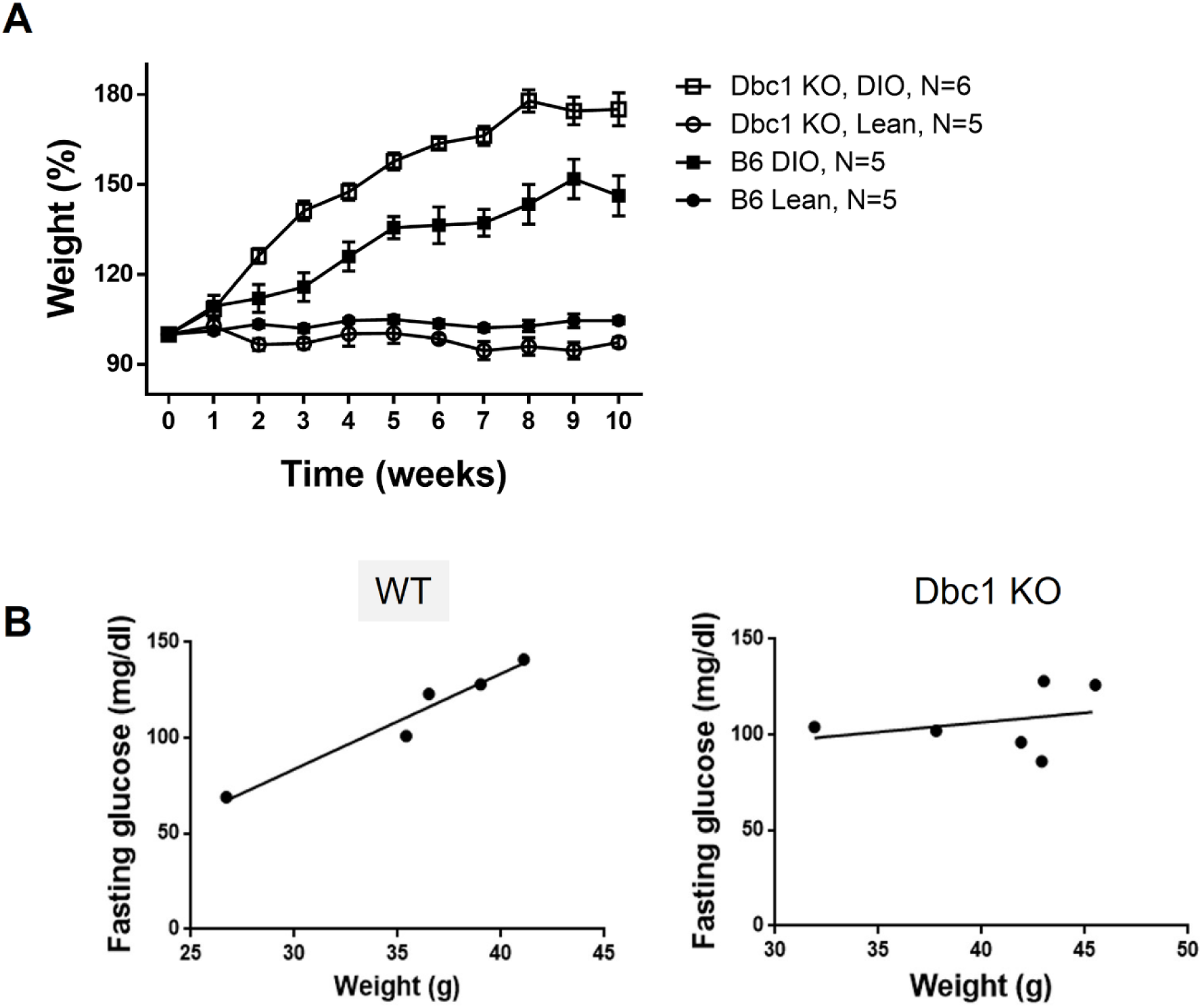
Metabolic protection of Dbc1 KO obese mice **(B)** Weight gain in WT and Dbc1 KO mice fed a high-fat diet on a C57BL/6J background, showing increased weight gain in Dbc1 KO mice compared to WT control mice. (**B)** Fasting glucose vs. body weight in obese mice shows a positive correlation between fasting glucose and body weight in WT, as expected. However, Dbc1 KO do not show this correlation, reflecting the protection against developing metabolic syndrome observed in Dbc1 KO animals.

**Figure S3.**
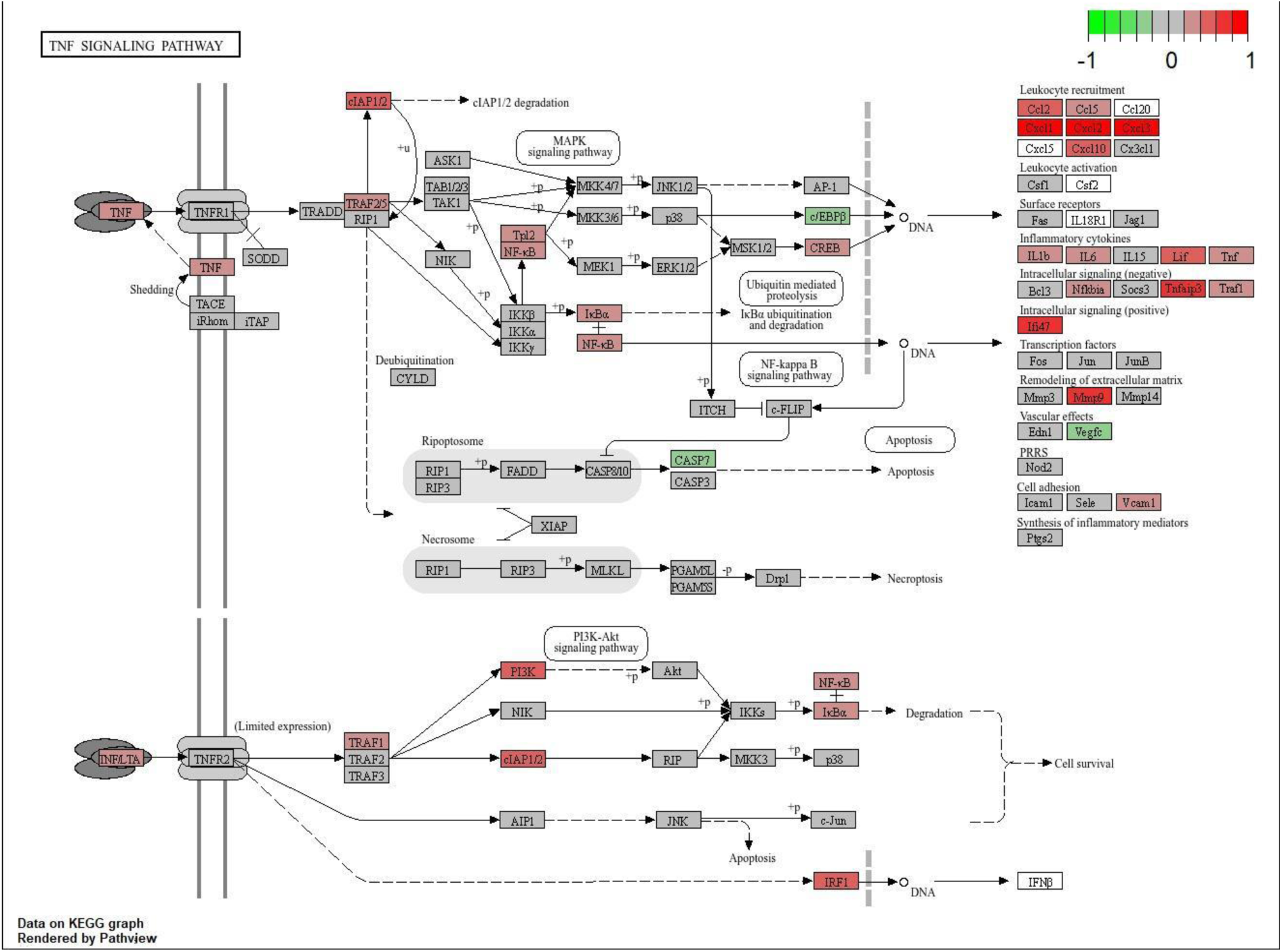
Pathway Analysis of Gene Expression Changes in Dbc1 KO adipocytes. The selected pathway, as diagrammed in KEGG, with gene expression changes (fold change, FC) projected based on the data. Genes marked in red are overexpressed in *Dbc1 KO*, while those in green are downregulated. Genes shown in gray indicate no change in expression between the two conditions.

**Figure S4.**
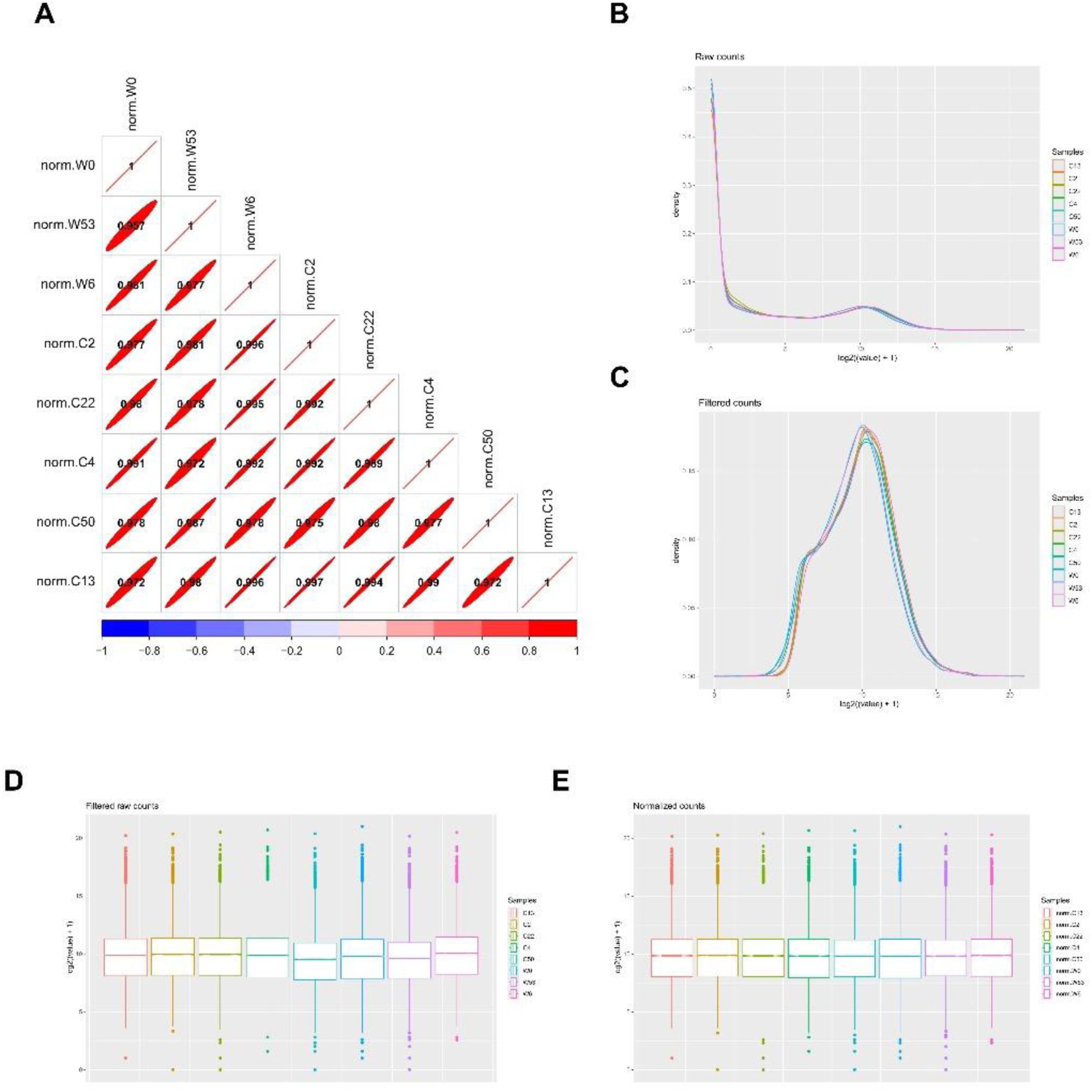
Analysis and processing of the RNA-seq samples (A) Pearson correlation matrix of the samples. Density plot of the samples pre (B) and post(C) filtering low read counts features. Boxplot of the samples pre (D) and post (E) normalization was applied

## References

1. Longo M, Zatterale F, Naderi J, Parrillo L, Formisano P, Raciti GA, et al. Adipose Tissue Dysfunction as Determinant of Obesity-Associated Metabolic Complications. Int J Mol Sci. 2019 Jan;20(9):2358.

2. Escande C, Nin V, Pirtskhalava T, Chini CC, Thereza Barbosa M, Mathison A, et al. Deleted in Breast Cancer 1 regulates cellular senescence during obesity. Aging Cell. 2014 Oct;13(5):951–3.

3. Escande C, Nin V, Pirtskhalava T, Chini CCS, Tchkonia T, Kirkland JL, et al. Deleted in breast cancer 1 limits adipose tissue fat accumulation and plays a key role in the development of metabolic syndrome phenotype. Diabetes. 2015 Jan;64(1):12–22.

4. Trauernicht AM, Kim SJ, Kim NH, Boyer TG. Modulation of estrogen receptor alpha protein level and survival function by DBC-1. Mol Endocrinol Baltim Md. 2007 Jul;21(7):1526–36.

5. Fu J, Jiang J, Li J, Wang S, Shi G, Feng Q, et al. Deleted in Breast Cancer 1, a Novel Androgen Receptor (AR) Coactivator That Promotes AR DNA-binding Activity. J Biol Chem. 2009 Mar 13;284(11):6832–40.

6. Li Z, Chen L, Kabra N, Wang C, Fang J, Chen J. Inhibition of SUV39H1 methyltransferase activity by DBC1. J Biol Chem. 2009 Apr 17;284(16):10361–6.

7. Chini CCS, Escande C, Nin V, Chini EN. HDAC3 Is Negatively Regulated by the Nuclear Protein DBC1. J Biol Chem. 2010 Dec 24;285(52):40830–7.

8. Hiraike H, Wada-Hiraike O, Nakagawa S, Koyama S, Miyamoto Y, Sone K, et al. Identification of DBC1 as a transcriptional repressor for BRCA1. Br J Cancer. 2010 Mar 16;102(6):1061–7.

9. Park SH, Riley P, Frisch SM. Regulation of anoikis by Deleted in Breast Cancer-1 (DBC1) through NF-κB. Apoptosis Int J Program Cell Death. 2013 Aug;18(8):949–62.

10. Qin B, Minter-Dykhouse K, Yu J, Zhang J, Liu T, Zhang H, et al. DBC1 functions as a tumor suppressor by regulating p53 stability. Cell Rep. 2015 Mar 3;10(8):1324–34.

11. Committee for the update of the guide for the care and use of laboratory animals. Guide for the care and use of laboratory animals. National Academies Press; 2011.

12. Meikle MN, Schlapp G, Menchaca A, Crispo M. Minimum volume Spatula MVD vitrification method improves embryo survival compared to traditional slow freezing, both for in vivo and in vitro produced mice embryos. Cryobiology. 2018;84:77–81.

13. Schlapp G, Goyeneche L, Fernández G, Menchaca A, Crispo M. Administration of the nonsteroidal anti-inflammatory drug tolfenamic acid at embryo transfer improves maintenance of pregnancy and embryo survival in recipient mice. J Assist Reprod Genet. 2015 Feb;32(2):271–5.

14. Nakagata N. Reproductive Engineering Techniques in Mice. Technical manual. In p. 6– 13.

15. Escande C, Chini CCS, Nin V, Dykhouse KM, Novak CM, Levine J, et al. Deleted in breast cancer-1 regulates SIRT1 activity and contributes to high-fat diet-induced liver steatosis in mice. J Clin Invest. 2010 Feb;120(2):545–58.

16. Andrés-León E, Núñez-Torres R, Rojas AM. miARma-Seq: a comprehensive tool for miRNA, mRNA and circRNA analysis. Sci Rep. 2016 May 11;6(1):25749.

17. Martin M. Cutadapt removes adapter sequences from high-throughput sequencing reads. EMBnet.journal. 2011 May 2;17(1):10–2.

18. Kim D, Paggi JM, Park C, Bennett C, Salzberg SL. Graph-based genome alignment and genotyping with HISAT2 and HISAT-genotype. Nat Biotechnol. 2019 Aug;37(8):907–15.

19. Liao Y, Smyth GK, Shi W. featureCounts: an efficient general purpose program for assigning sequence reads to genomic features. Bioinformatics. 2014 Apr 1;30(7):923–30.

20. Varet H, Brillet-Guéguen L, Coppée JY, Dillies MA. SARTools: A DESeq2- and EdgeR-Based R Pipeline for Comprehensive Differential Analysis of RNA-Seq Data. PLOS ONE. 2016 Jun 9;11(6):e0157022.

21. Robinson MD, McCarthy DJ, Smyth GK. edgeR: a Bioconductor package for differential expression analysis of digital gene expression data. Bioinformatics. 2010 Jan 1;26(1):139– 40.

22. Wu T, Hu E, Xu S, Chen M, Guo P, Dai Z, et al. clusterProfiler 4.0: A universal enrichment tool for interpreting omics data. The Innovation. 2021 Aug 28;2(3):100141.

23. Nin V, Chini CCS, Escande C, Capellini V, Chini EN. Deleted in breast cancer 1 (DBC1) protein regulates hepatic gluconeogenesis. J Biol Chem. 2014 Feb 28;289(9):5518–27.

24. Chen EY, Tan CM, Kou Y, Duan Q, Wang Z, Meirelles GV, et al. Enrichr: interactive and collaborative HTML5 gene list enrichment analysis tool. BMC Bioinformatics. 2013 Apr 15;14(1):128.

25. Kong S, Thiruppathi M, Qiu Q, Lin Z, Dong H, Chini EN, et al. DBC1 is a suppressor of B cell activation by negatively regulating alternative NF-κB transcriptional activity. J Immunol Baltim Md 1950. 2014 Dec 1;193(11):5515–24.

26. Wernstedt Asterholm I, Tao C, Morley TS, Wang QA, Delgado-Lopez F, Wang ZV, et al. Adipocyte Inflammation Is Essential for Healthy Adipose Tissue Expansion and Remodeling. Cell Metab. 2014 Jul 1;20(1):103–18.

